# A Facile and Rapid Route to Self-Digitization of Samples into a High Density Microwell Array for Digital Bioassays

**DOI:** 10.1101/2021.01.31.428989

**Authors:** Xu Cui, Tianbao Hu, Qiang Chen, Qiang Zhao, Yin Wu, Tengbao Xie, Pengyong Liu, Xi Su, Gang Li

## Abstract

Digital bioassays are powerful methods to detect rare analytes from complex mixtures and study the temporal processes of individual entities within biological systems. In digital bioassays, a crucial first step is the discretization of samples into a large number of identical independent partitions. Here, we developed a rapid and facile sample partitioning method for versatile digital bioassays. This method is based on a detachable self-digitization (DSD) chip which couples a reversible assembly configuration and a predegassing-based self-pumping mechanism to achieve an easy, fast and large-scale sample partitioning. The DSD chip consists of a channel layer used for loading sample and a microwell layer used for holding the sample partitions. Benefitting from its detachability, the chip avoids a lengthy oil flushing process used to remove the excess sample in loading channels and can compartmentalize a sample into more than 100,000 wells of picoliter volume with densities up to 14,000 wells/cm^2^ in less than 30 s. We also demonstrated the utility of the proposed method by applying it to digital PCR and digital microbial assays.

---

Recently digital assays have rapidly emerged as key technologies for the detection or analysis of single biological entities such as molecules, organisms or cells.^1–4^ In such assays, a sample is divided into a large number of equal volume subunits to isolate single entities, namely, “digitization”. This “digitization” endows digital assays with a number of outstanding abilities, including: i) absolutely quantifying biomolecules without reference standards;^5–7^ ii) measuring single-molecule kinetics;^8–10^ iii) probing single-cell phenotype and genotype;^11^ iv) studying temporal processes of chemical or biochemical reactions on microscales.^12–14^ These abilities prompt the widespread application of digital assays, especially in the detection of very minute quantities of diagnostic markers, quantitative assays for DNA and protein markers.^15–18^ Moreover, digital assays are also finding use in quantitative identification of microbials,^19–21^ investigation of nucleation kinetics,^12–14^ measurement of single enzyme kinetics,^8–10^ and dissection of cellular heterogeneity.^22–24^

In digital assays, “digitization” is crucial, which affects the dynamic range, precision and robustness of assays. To date, various sample “digitization” methods have been developed for the application of digital assays in industry and academic research.^2, 4^ Among them, droplet microfluidics is the most popular method to generate a large number of droplets for digital assays. In general, monodisperse aqueous droplets are generated in an oil phase using microfluidic T-junction, flow-focusing or step geometries.^5, 6, 8, 12–14, 19, 23–27^ However, most of droplet microfluidic platforms require external equipment. In addition, monodisperse droplet generation and stabilization require the addition of surfactants, which may have some impacts on the feasibility of bioassays.^28^ Another disadvantage of droplet microfluidic method is the difficulty in time-lapse monitoring of large numbers of individual drops since droplets are free to move throughout the experiment, making them unsuitable for studying temporal processes. To address these issues, an alternative strategy is to partition a sample into physically isolated compartments using pneumatic valve arrays,^29^ slipchip,^30^ high-density thru-holes,^31, 32^ interfacial tension-assisted trapping.^33–36^ Because all partitions are physically confined inside chambers or holes and there are solid barriers between adjacent partitions, the compartment-based methods avoid the risk of droplet coalescence during reaction or incubation, thus ensuring the accuracy of measurement or detection. Moreover, the immobilized array format of the compartment-based partitioning also offers remarkable benefits for studying temporal processes of large numbers of individual entities since the partitions can be spatially indexed and be time-lapse monitored. Despite their obvious advantages, the conventional compartment-based methods still face a number of challenges, such as requirement of additional mechanical components or expensive equipment, lengthy partitioning process, and costly consumables (chips or specific oil). To simplify the fluid operation and reduce the assay cost, a predegassing-based self-pumping mechanism was introduced in several compartment-based platforms for partitioning sample automatically, with no external equipment and minimal manual intervention.^37–42^ These predegassing-driven self-digitization (SD) platforms obviate the external control instruments and make digital assays more accessible to common researchers. However, these SD platforms generally require the removal of the excess sample in loading channel using oil flushing to isolate all partitions. To achieve effective confinement of the partitions and robust prevention of crosstalk between partitions, high viscosity or thermo-curing oils are generally used for oil flushing, which results in a lengthy partitioning process, thus making these SD platforms unfavorable to most of applications.

To address this challenge, we present a new SD strategy for rapid, simple and low-cost digital assays. Unlike the existing predegassing-driven SD methods, this strategy combines a reversibly assembled chip with the predegassing-based self-pumping mechanism. In the reversibly assembled chip, the sample loading channels and partitions holding wells are located on separate layers; we call the device a detachable self-digitization (DSD) chip to distinguish it from the monolithic channel-well configuration reported previously. ^36–39, 41, 42^ As schematically depicted in Figure 1, the sample loading function layer contains an array of parallel microfluidic channels which are all connected to sample inlet *via* four main channels, and the partitions holding function layer contains a high-density array of microwells. In the state of assembly, the sample loading channels overlap the microwell array, ensuring the priming of all microwells during sample loading. Benefitting from its reversible assembly, the chip can rapidly partition a sample into arrays of independent wells by directly detaching the loading channel layer, without the requirement of a lengthy oil flushing. Furthermore, the separate configuration of the sample loading function and partitions holding function in the DSD chip allows more available space for partitioning wells, which is favorable for wide dynamic range and high precision detection. Based on this strategy, we developed a glass slide-sized DSD chip which can facilely and uniformly partition biological entities in an array of more than 100,000 wells with densities up to 14,000 wells/cm^2^ in less than 30 s. Finally, we also demonstrated the capability of the proposed DSD chip by applying it to digital PCR and digital microbial assays.

**Figure 1.**
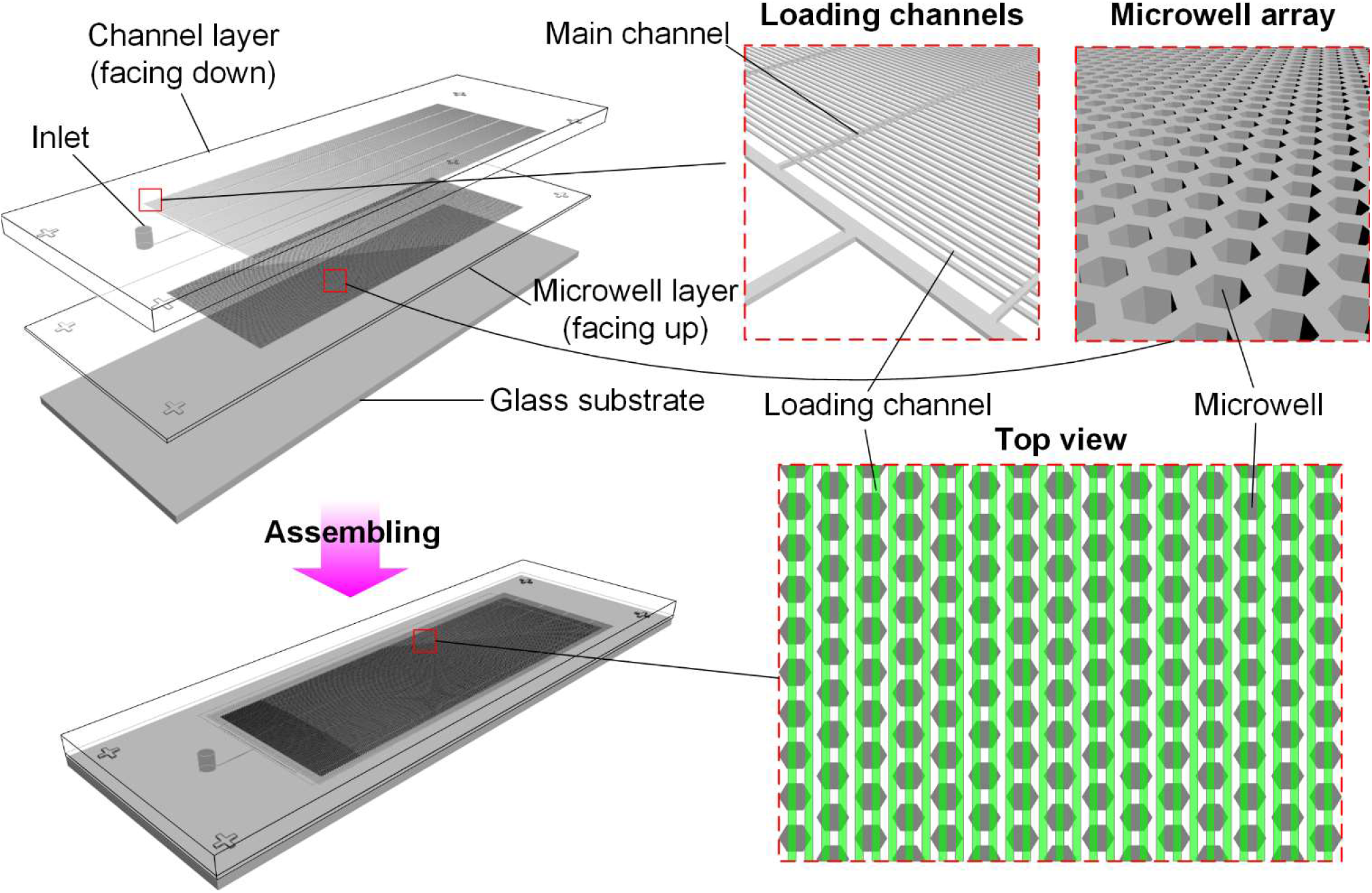
Schematic of the detachable self-digitization chip.

## EXPERIMENTAL SECTION

### Chip Design and Fabrication

Figure 1 shows the schematic of the DSD chip, which is reversibly assembled from three layers: a PDMS channel layer, a PDMS microwell layer, and a supporting glass substrate. The channel layer contains an array of 667 parallel dead-end microfluidic channels used for loading sample, and one end of each channel connects to a main channel which connects to an inlet port. The microwell layer contains an array of 113,140 microwells used to hold the sample partitions. To facilitate the alignment of the loading channels to the partition microwells, the footprint of the loading channel array is designed to be 1.2 mm offset outside the footprint area of the microwell array. In addition, the gap of neighbor loading channels is smaller than the maximum width of the microwell, which ensures each microwell partly overlaped with at least one channel, and then achieving a complete priming of all microwells. The chip has the same footprint as a standard microscope slide (25 mm × 75 mm), making it compatible with various existing microscope systems and microarray scanners.

The proposed DSD chip was fabricated mainly based on the multi-layer soft lithography as described previously. Briefly, using standard lithographic procedures, SU-8 3010 and 3050 negative photoresist (MicroChem Corp., Newton, MA) were patterned onto separate silicon wafers to create two molds for the channel layer and microwell layer, respectively. Typical dimensions of the molds are: main channel = 52 mm (length) × 80 μm (width) × 60 μm (height); loading microchannel = 17.9 mm (length) × 30 μm (width) × 20 μm (height); gap of neighbor channels = 48 μm; microwell = 70 μm (diagonal) × 40 μm (height). Prior to PDMS casting, the prepared SU-8 molds were silanized with trichloro(1H, 1H, 2H, 2H-perfluorooctyl) silane (Sigma-Aldrich China, Shanghai, China) to facilitate the release of PDMS replicas. Then, according to different applications, PDMS prepolymer was prepared with two different compositions: i) PDMS monomer and cross-linker (Sylgard 184, Dow Corning, Midland, MI) were mixed with a 10:1 weight ratio to fabricate DSD chips for digital microbial assays; and ii) PDMS monomer, cross-linker and Triton X-100 (Sigma-Aldrich China, Shanghai, China) was mixed with a 10:1:0.05 weight ratio to fabricate DSD chips for digital PCR. After degassing for 1 hr, the PDMS prepolymer were poured over the molds with different thickness: 0.5 mm for the channel layer and 2 mm for the microwell layer. After curing at 70 °C for 3 hr, the PDMS replicas were cut and peeled off from the molds and an inlet port was created on the PDMS channel layer with a punching tool. Next, the unstructured surface of the PDMS microwell layer was bonded to a glass slide by plasma treatment. Finally, the PDMS channel layer and the PDMS microwell layer were face-to-face aligned and conformally contacted to form a reversible seal.

### Chip Operation

The basic operation of the DSD chip is outlined in Figure 2. Firstly, the inlet and outlet of the chip was sealed with tape, and then the sealed chip was degassed in a desiccator (~ 10 kPa) for 1 hr to build up a vacuum in microchannels and microwells for self-priming. After removing the chip from the desiccator, a pipette tip holding 20 μL of sample was immediately inserted through the tape and into the inlet of the chip. About 5 s, all of channels and microwells were completely filled with the sample, and then the channel layer was peeled off from the microwell layer and an accompanying oil sealing was applied to isolate the sample in each microwell. Corresponding to different applications, the well layer containing the partitioned sample can be sealed in two different modes: i) sandwich mode, *i.e.*, the well layer was sealed by sandwiching it with a liquid-PDMS-coated glass; ii) oil-covering mode, *i.e.*, the well layer was directly immersed into oil. Finally, the chip sealed with sandwich mode was placed on a flat PCR apparatus for amplification reaction and the chip sealed with oil-covering mode was fixed on a microscope stage for monitoring the growth of bacteria.

**Figure 2.**
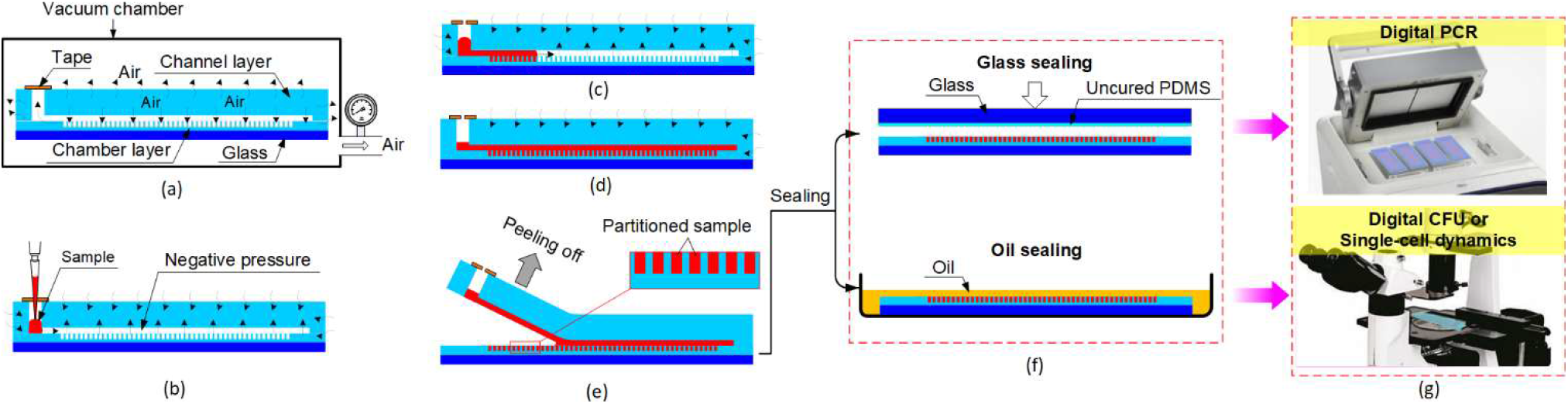
Schematic illustration of the operation procedure of the DSD chip. (a) degassing of the chip in a vacuum chamber; (b) removing the chip from vacuum conditions and adding an aliquot of sample in the inlet; (c) aspiration of the sample into the microchannels under the negative pressure created by the degassed PDMS substrate; (d) sample priming of all channels and wells; (e) peeling off the channel layer to isolate all the primed wells; (f) sealing of the partitioned sample with a PDMS-coated glass slide or a layer of oil; (g) transferring of the DSD chip on a flat PCR instrument for digital PCR assay or on a stage of microscope for time-lapse monitoring of single biological entities.

### PCR amplification

In this study, BRAF-V600E mutation detection was performed to demonstrate the feasibility of our DSD chip for digital PCR. BRAF-V600E genomic DNA was extracted from the RKO cell line using the animal tissue/cell genomic DNA extraction kit (Solarbio, Beijing, China) and DNA concentrations were calibrated with a NanoDrop spectrometer (ThermoFisher Scientific, Massachusetts, USA). A dilution series of target DNA was prepared using Tris-EDTA buffer. The total reaction volume of each digital PCR was 20 μL, which contained 2 μL 10×dPCR Buffer (Turtle Tech, China), 0.57 μL DNA polymerase, 0.8 μL forward primer (10 μM), 0.8 μL reverse primer (10 μM), 0.8 μL probe (10 μM), 14.03 μL RNase-free water and 1 μL target template diluted by Tris-EDTA buffer. All primers and probes for digital PCR were purchased from Turtle Tech (Shanghai, China). The sequences are given as follows (all in 5′ → 3′ direction): Forward primer: CACCTCAGATATATTTCTTCATG; Reverse primer: TTCAAACTGATGGGACCC; Probe: [FAM]-AGA-TTTCTCTGTAGC-[MGB]. Before being loaded in the DSD chip, all reaction components were premixed off-chip. After the completion of sample self-digitization and chip sealing, the chips were placed on a thermocycler (MasterCycler Nexus flat, Eppendorf) to perform PCR thermocycling. The thermocycling protocol consisted of 50 °C for 10 min, 95 °C for 10 min, and 45 cycles of 45 s at 95 °C and 40 s at 60 °C. Sterile double-distilled water without template was used as negative control in each run of dPCR. All dPCR experiments were performed in triplicate.

### Counting of Viable Bacteria

In this study, a fluorescent strain of Escherichia coli (*E. coli* BL21) was used to demonstrate the feasibility of our DSD chip for the precise counting of viable bacteria. Firstly, single colonies of *E. coli* BL21 strain were prepared by reviving a frozen glycerol stock and streaking a plate (see section S1 in Supporting Information for more details). Then, an isolated single colony of *E. coli* BL21 from the plate was inoculated in LB broth (Land Bridge, Beijing, China) and pre-cultured for 6 hr (37°C with a shaking rate of 220 rpm) to obtain *E. coli* BL21 stock with concentration of ~ 10^9^ cells/mL. Then, a conventional plating-based quantification assay was carried out to measure the concentration of *E. coli* BL21 stock (see section S2 and Figure S1 in Supporting Information for more details). After that, the *E. coli* stock was serially diluted with LB broth and the obtained sample concentrations ranged from 10^2^ to 10^7^ cells/mL. Then, the diluted *E. coli* samples were loaded in six predegassed DSD chips respectively for partitioning. After partitioning and oil-sealing, the chips were incubated at 37 °C for 5 hr in an incubator (LHP-160, Jiebosen Instrument, Changzhou, China). Finally, the chips were taken out from the incubator and microscopically examined.

### Monitoring of single bacteria growth kinetics

Dynamic measurements of single bacteria growth kinetics were also made on the *E. coli* BL21 strain. Firstly, an isolated single colony of *E. coli* BL21 was pre-cultured in LB broth for 6 hr (37°C with a shaking rate of 220 rpm) to obtain a cell concentration of ~10^9^ CFU/mL. After that, the *E. coli* sample was diluted 10^6^ times with LB broth, and then loaded in the predegassed SD chip for partitioning. After partitioning and oil-sealing, the chip was placed in an incubator (LHP-160, Jiebosen, Changzhou, China) for incubation at 37°C. The colonial growth of single bacteria was examined by fluorescence microscopy at regular intervals of 2 hr.

### Chip Imaging and Data Analysis

In this study, all bright-field and fluorescence images were taken with an upright microscope (CX40, Sunny Optical, China) which was equipped with a mercury lamp and a CMOS camera (SOPTOP OD230R, Sunny Optical, China). ImageJ software (http://rsbweb.nih.gov) was used to analyze the obtained fluorescence images. In the case of digital PCR, the number of positive wells in each SD chip were counted with ImageJ, and then the average copy number of template in each SD chip were determined by combining the counting data with Poisson statistics principle. In the case of digital cell assays, the total fluorescence intensity of each well containing microcolony in the SD chips was measured with ImageJ (see section S3 in Supporting Information for more details). Next, the obtained data were input into the USDA integrated pathogen modeling program (IPMP 2013, https://www.ars.usda.gov/) for modeling of the bacterial growth curve. Finally, OriginPro 2018 software (OriginLab, Northampton, MA) was used for further data processing.

## RESULTS AND DISCUSSION

### Facile and Rapid Sample Self-Digitization

The DSD chip is designed to simplify and accelerate digital assays. To this end, two critical technologies were integrated in the DSD chip: i) predegassing-based vacuum storage in PDMS, and ii) detachable channel-well assembly. Figure 2 shows the straightforward user protocol for sample self-digitization with the DSD chip, which only involves three steps: i) chip degassing, ii) sample loading/priming, and iii) wells isolation. Incidentally, the overall hand-on time of sample self-digitization can be greatly reduced because the DSD chip can be degassed in advance and then stored in airtight packages (such as aluminium foil pouches) for ready use. To demonstrate the applicability of the DSD chip to digital bioassays, we first tested its capability of sample self-digitization. To allow easy visualization, a red food dye was used as a sample for self-digitization test. Driven by the negative pressure in the microchanel/microwells built up during degassing, the dye solution pipetted in the inlet port of the DSD chip was automatically sucked into the main channel and quickly filled all of the available space in the chip including loading channels and partitioning wells (Figure 3 and Movie S1 in Supporting Information). The well array was completely primed with sample in less than 5 s. After the completion of priming, the channel layer was removed from the well array layer to immediately isolate all primed wells and achieve a rapid partitioning. Note that in contrast to the previous compartment-based digital platforms which were in monolithic channel-well configurations,^36–39, 41, 42^ our DSD chip had a separated channel-well configuration. This configuration allows a direct removal of the excess sample interconnecting the primed wells *via* detachment without a lengthy flushing step, and thus resulting in an ultra-fast sample self-digitization. This self-digitization method can prime and partition an aqueous sample into 113140 wells in less than 30 s, so its discretization rate is equivalent to generation of droplets with frequencies of ∼ 3.8 kHz in droplet microfluidic devices. Furthermore, this separate configuration offers three other advantages: i) allowing dead-end and outlet-free channels for sample loading which is favorable for lowering sample consumption; ii) allowing denser well arrays per chip area which is favorable for wide dynamic range and high precision detection; iii) allowing to shift to an open configuration which is favorable for the retrieval of specific partitions and downstream biological analysis. As shown in Figure 3b, a uniform and defect-free partitioning was achieved with the DSD chip. All the wells (100%) were filled up with the solutions.

**Figure 3.**
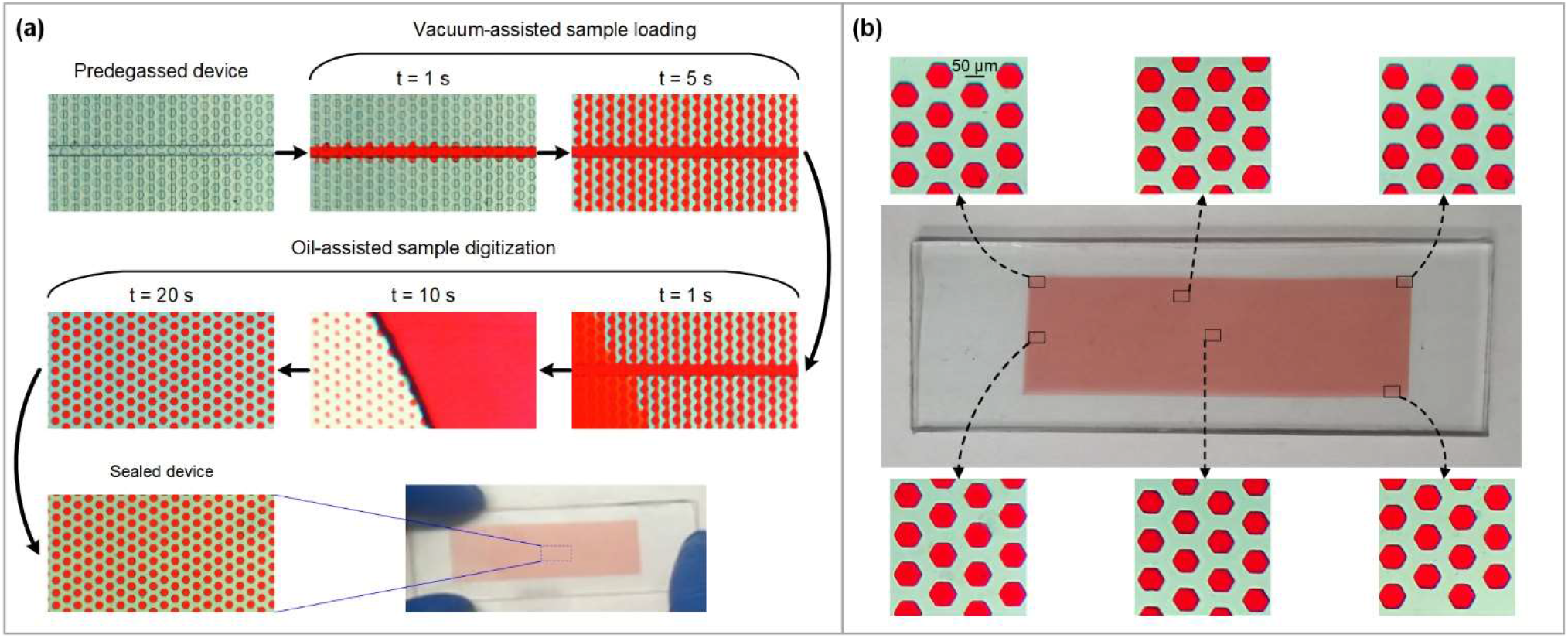
Illustration of the DSD chip for rapid and uniform sample digitization. (a) Time-lapse of the sample digitization process. (b) Microscopic images of the partitioned droplets from six random positions on the DSD chip.

### Absolute Quantification of Target DNA

Detection of BRAF mutations is important for the clinical management of colorectal carcinomas (CRC). Particularly, the determination of BRAF-V600E mutation, a predominant BRAF mutation, provides a diagnostic exclusion criterion for hereditary nonpolyposis colorectal cancer. To explore the capability of our DSD platform, we firstly applied it to quantitative digital PCR detection of BRAF-V600E mutation. A series of DNA template solutions was prepared with concentrations ranging from 10 copies/μL to 10^5^ copies/μL. As described in the section of “Chip Operation”, six premixed PCR reactions, including the negative control, were loaded into six predegassed DSD chips, respectively, and partitioned into a large number of microwells. After partitioning, we assembled the DSD chip with a glass-PDMS-glass sandwich configuration (as shown in Figure 2), which can prevent the reagent evaporation in the chip during thermal cycling. Subsequently PCR thermocycling and fluorescent imaging were performed. Figure 4a shows representative fluorescence images obtained from different concentrations of DNA template. The obtained fluorescence images were then analyzed to determine the percentage of positive partitions. Finally, the average copy number of DNA template in each chip was calculated according to the Poisson distribution. The results showed that there was a good linear correlation between the measured template concentrations and their expected values (R^2^ = 0.9998), as shown in Figure 4b. Furthermore, the slope and intercept of the linear correlation were measured as 1.0021 and 0.2964, respectively, demonstrating the measured concentrations agree well with the expected concentrations. Note that that our DSD device was able to detect template DNA at concentrations as low as 10 copies/μL, with a dynamic range of over 5 orders of magnitude, demonstrating that the DSD chip is a useful platform for highly sensitive and precise genetic analysis.

**Figure 4.**
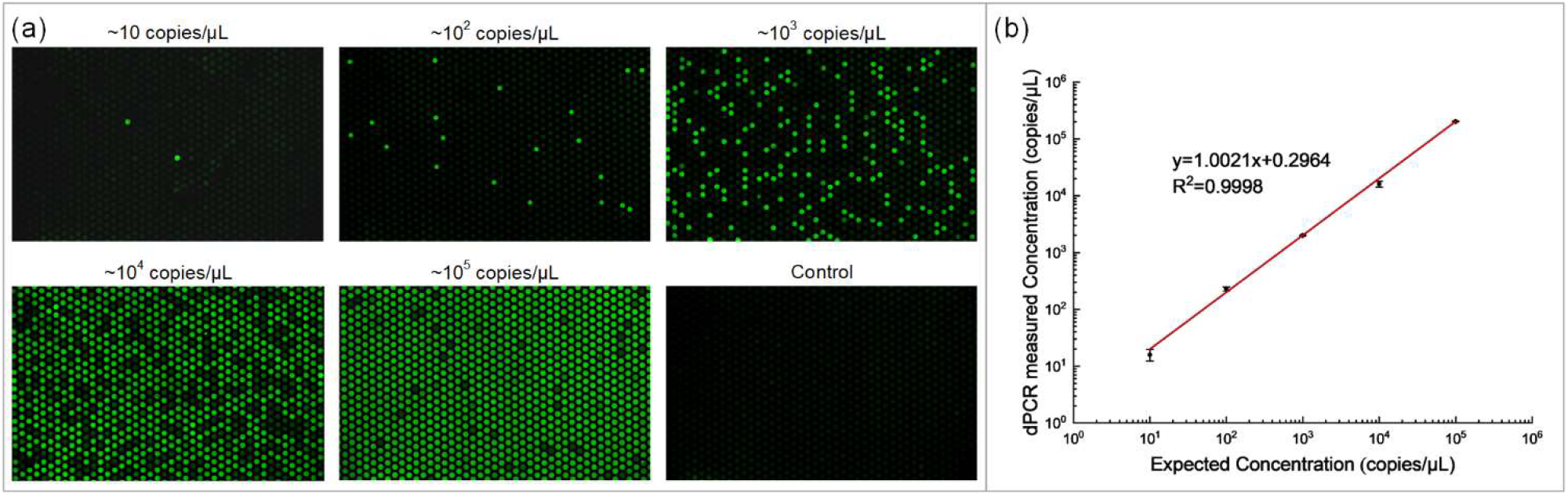
Digital PCR on the DSD chip with different concentration of BRAF-V600E genomic DNA: (A) Digital PCR on the DSD chip with a serial dilution of target DNA template ranging from 10 copies/μL to 10^5^ copies/μL. The negative assay was in control when no target template was loaded. (B) The linear relationship between the measure value (copies/μL) in the DSD chip and the expected copy number per reaction. The measure values matched well with the expected values according to Poisson statistics (R^2^=0.9998).

### Precise Enumeration of Bacteria over a Dynamic Range

A precise enumeration of viable bacteria is important to a variety of microbiological applications such as clinical diagnosis, food quality monitoring and antibiotic susceptibility testing. Traditionally, plate counting and measurement of turbidity (OD) have been used for quantifying viable bacteria in samples, but they have some limitations, such as cumbersome manual operation, labor-intensive analysis, long incubation time, or relatively inaccurate enumeration. Here, we demonstrate that the DSD chip can be used to simply, rapidly and precisely quantify viable bacteria over a wide dynamic range of colony forming unit (CFU) densities. By stochastically isolating bacteria into a large number of microwells, a precise quantification of bacteria can be achieved. Assuming that bacteria are homogeneously suspended in the original sample solution and the partitioned droplets in microwells are monodisperse, the distribution of the bacterial number per microwell follows Poisson law. Therefore, based on the Poisson theory, the proportion of positive microwells containing at least one bacterium can be used to calculate the initial bacterial concentration in the sample. This microbiological quantification method is similar to digital PCR in quantitative strategy and thus can be named as “digital CFU” (dCFU). Although single bacteria can be observed under the microscope at a 400 × magnification without waiting for the colonies to grow, it is impractical for this examination mode to directly count the number of positive microwells containing bacteria in the chip due to the small size of individual bacteria and narrow field of view under high microscopic magnification. Fortunately, the colony growth of single bacteria can facilitate the bacterial detection. Generally, after incubation of 4 ~ 5 hr, the colony becomes visible under a low magnification (*e.g.* 100 ×), and then the number of positive microwells containing colonies in a chip can be counted, which is much faster than the traditional plating method (typically requiring 12 ~ 24 hr). As a proof of concept, we used the DSD chip-based dCFU method to quantify the viable bacteria in samples. Similar to digital PCR, a series of expected concentrations of *E. coli* BL21 were loaded and partitioned in DSD chips for dCFU counting. After 5 h incubation at 37 °C, all chips were examined by fluorescent microscopy. Figure 5a shows representative fluorescence images obtained from different concentrations of *E. coli* BL21. Meanwhile, to validate the quantitative ability of our dCFU method, we compared the results (*i.e.* calculated number of CFUs in the dilutions) obtained with our dCFU assay and the expected concentrations. As shown in Figure 5b, the *E. coli* BL21 concentrations measured with the DSD chip-based dCFU correspond very well to the expected concentrations (R^2^ = 0.9975) in the entire 5 log range of the measured sample concentrations, demonstrating the excellent reliability of the DSD chip-based dCFU method for the quantification of microbes. Compared to the conventional plate counting method, the DSD chip-based dCFU method provides three distinct advantages: i) faster bacteria detection; ii) lower sample/reagent consumptions; iii) less labor-consuming; iv) wider dynamic range of concentrations. Furthermore, unlike the droplet-based dCFU method, in which high shear forces act on the bacteria during encapsulation of bacteria and reduce the bacterial viability,^20^ the DSD chip-based dCFU method loads little shear forces on the bacteria during sample partitioning, and thus the shear force-induced damage to bacteria can be ignored. By marrying the simplicity of plating and the speed of microscopy, this DSD chip-based method achieves a simple, fast, and precise counting of viable bacteria, providing an extremely helpful tool for various microbiological applications.

**Figure 5.**
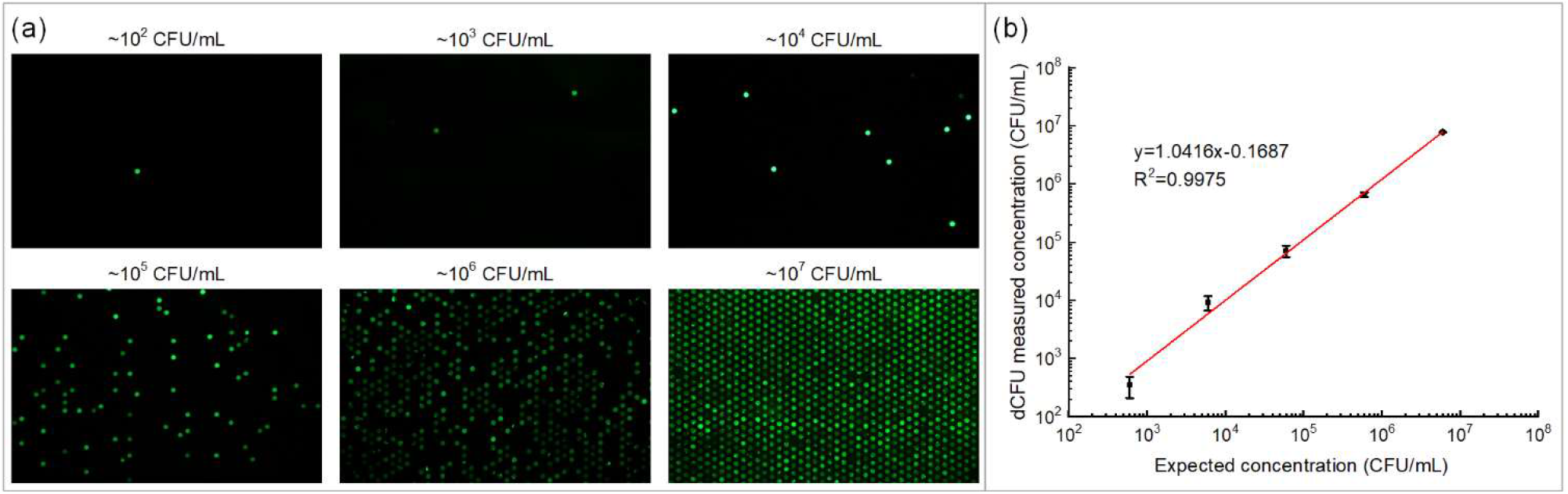
Digital enumeration of bacteria *via* the DSD chip. (a) Representative images of growth of different concentration fluorescent *E. coli* BL21 in the DSD chip. (b) The linear relationship between the measured concentrations with dCFU assay and the expected concentration of *E. coli* BL21 samples whose concentrations ranged over five orders of magnitude. Strong linearity between the calculated number of cells and the expected number of cells for *E. coli* BL21 indicate that DSD chip can be applied for the precise enumeration of bacteria in a wide dynamic range.

### Monitoring the Growth Kinetics of Single Bacteria

It is well known that population heterogeneity highly affects the behaviour of the microbial populations, including growth, survival, adaption, infection, and biofilm formation. However, conventional bacterial growth studies rely on cultivated bulk populations, in which cellular heterogeneity is typically fully masked by the average bulk behaviour. To reveal and identify the microbial population heterogeneity, investigation of the growth of bacteria in single-cell level is essential. One of the attractive features of our DSD chip is the ability to partition a bulk sample in a large number of microwells, forming a high-density stochastic confinement in small volumes, so that single bacteria can be isolated from the bulk mixture. Such stochastic confinement enables simultaneous monitoring of the growth of single bacteria encapsulated in individual droplets for heterogeneity analysis. As an example, we show how the DSD chip can be used to track the growth of *E. coli* BL21 single cells. Similar to the dCFU experiments mentioned above, a diluted bacteria solution (~ 10^3^ CFU/mL) was firstly loaded in a DSD chip and partitioned into a high density microwell array. Then, bacterial growth was observed through an increase in fluorescence intensity over time. As shown in Figure 6a, three fluorescent monoclonal colonies of *E. coli* BL21 cultivated in microwells were monitored every 2 hr. In view of a direct proportional relationship between the number of bacteria in each colony and its total fluorescence intensity, we measured the total fluorescence intensity in each microwell as a proxy for the total number of bacteria in the microwell, obtaining the growth curves shown in Figure 6b. It can be found that there is a similar growth trend for 3 microcolonies but a variability in the final colony size and the growth rate among 3 microcolonies, demonstrating the heterogeneity in the growth dynamics of microcolonies originating from single bacteria. To quantify this heterogeneity, we fitted the growth data of each microcolony to the Huang primary model for the estimation of the growth kinetic parameters (see Figure S2 in Supporting Information).^43^ These kinetic parameters include: i) the lag time, λ, which is the period during which there is no increase in number of cell due to physiological adaptation of cells to the new environment; ii) the maximum specific growth rate, μ_max_, which is the maximum value of the logarithmic growth rate and can be estimated by the slope of the tangent drawn to the inflection of the sigmoid curve; iii) the final colony size, y_max_, corresponding to the maximum value of the fluorescence intensity of a microcolony after it reached a stationary phase. The table beside Figure 6b shows the growth parameters estimated by the primary model for these three microcolonies originating from single cells; the values of λ, μ_max_ and y_max_ varied from 3.08 hr to 3.32 hr, from 6.42 hr^−1^ to 6.62 hr^−1^, and from 11.95 a.u. to 12.57 a.u., respectively. Although three cells came from the same seed culture and grown in identical conditions, the final cell numbers (or colony size) were significantly different, which can be inferred from the total fluorescence intensities of 3 microcolonies. In repeated experiments, we observed similar heterogeneous growth for single bacteria. Furthermore, tracking individual bacteria growth curves also can be used to determine correlations between the growth parameters, which is helpful for building models of bacterial growth.^44^

**Figure 6.**
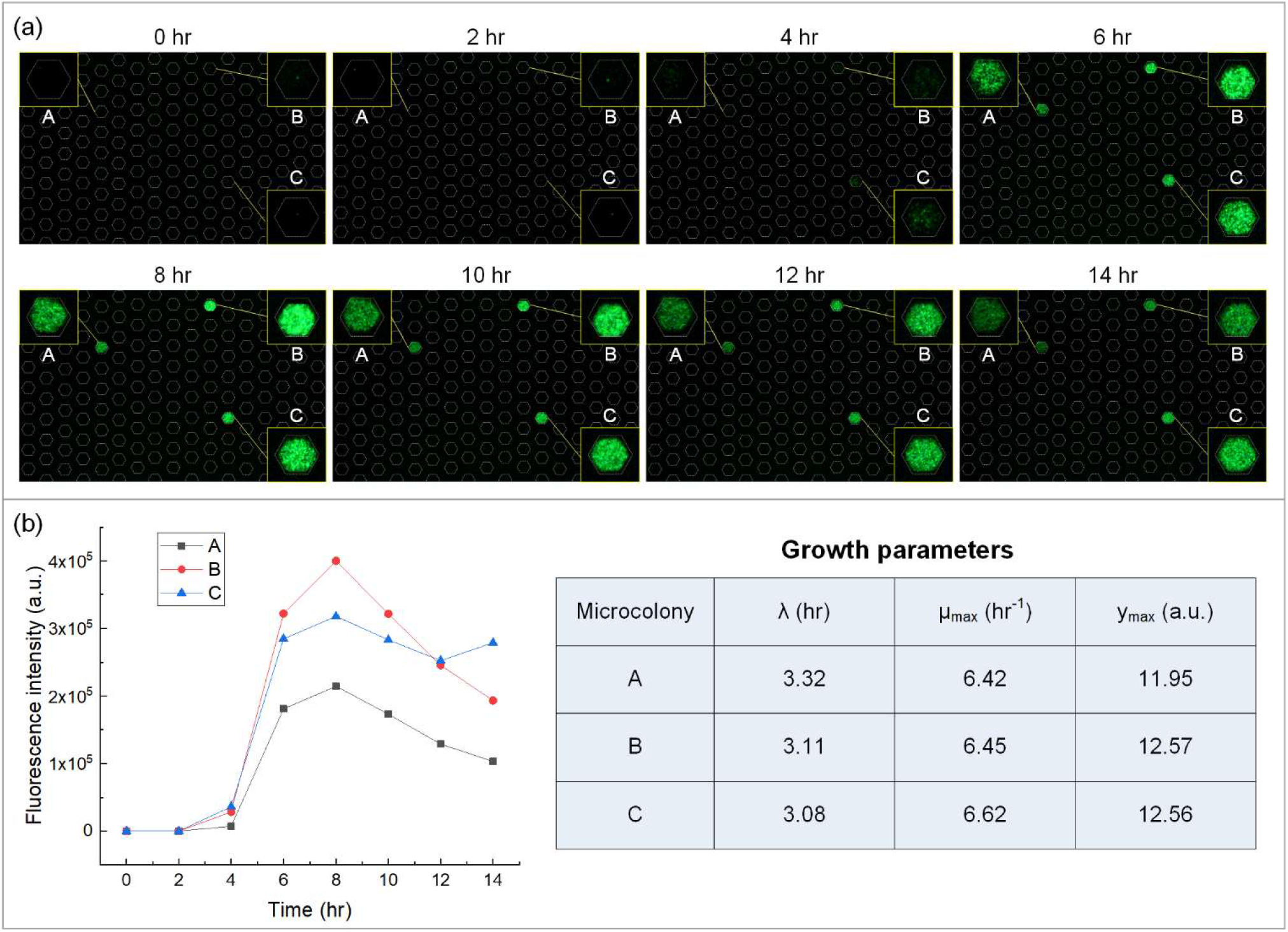
Single bacteria growth kinetics. (a) Time-lapse monitoring of growth of three *E. coli* BL21 colonies originated from single cells. (b) Growth curve of three *E. coli* BL21 colonies, the table beside shows the lag time, (λ), the maximum specific growth rate (μ_max_), and the final colony size (y_max_).

## CONCLUSION

In summary, we have developed a facile sample digitization method that can discretize a sample into more than 100,000 picoliter-sized subunits in less than 30 s. Based on a reversibly assembled microwell array device combining with a degas-created self-pumping mechanism, this method provides a simple, robust and versatile tool for digital bioassays. Unlike the existing microwell-based digital assay platforms where the well array and loading channels share a structure layer, this device divides the well array and loading channels into two different layers and both layers are reversibly assembled. Such features provide several advantages for digital assays: i) higher well densities for wide dynamic range assays, ii) straightforward and rapid isolation of all wells without a lengthy oil flushing, iii) greater sample utilization without filling waste or outlet ports during the sample loading. We demonstrated the capacities of this method by applying it to absolute quantification of target DNA (digital PCR), precise enumeration of bacteria (digital CFU), and time-resolved monitoring of the growth kinetics of single bacteria. The proposed method is technically simple and offers a route to reduce the cost of digital assays and to increase their accessibility to biology labs. We envision that this platform will be a powerful and cost-effective tool for digital bioassays.

## AUTHOR INFORMATION

### Author Contributions

The manuscript was written through contributions of all authors. / All authors have given approval to the final version of the manuscript. /

### Notes

The authors declare no competing financial interest.

## ACKNOWLEDGMENT

This work was supported by grants from the National Natural Science Foundation of China (No. 61974012, 21827812, 52005063 and 61771078), the Chongqing Research Program of Basic Research and Frontier Technology (No. cstc2017jcyjBX0036), the Natural Science Foundation of Chongqing (No. cstc2020jcyj-msxmX0188), the National Postdoctoral Program for Innovative Talents (No. BX20190049), and the Graduate Scientific Research and Innovation Foundation of Chongqing (No. CYB19035).

---

For TOC only

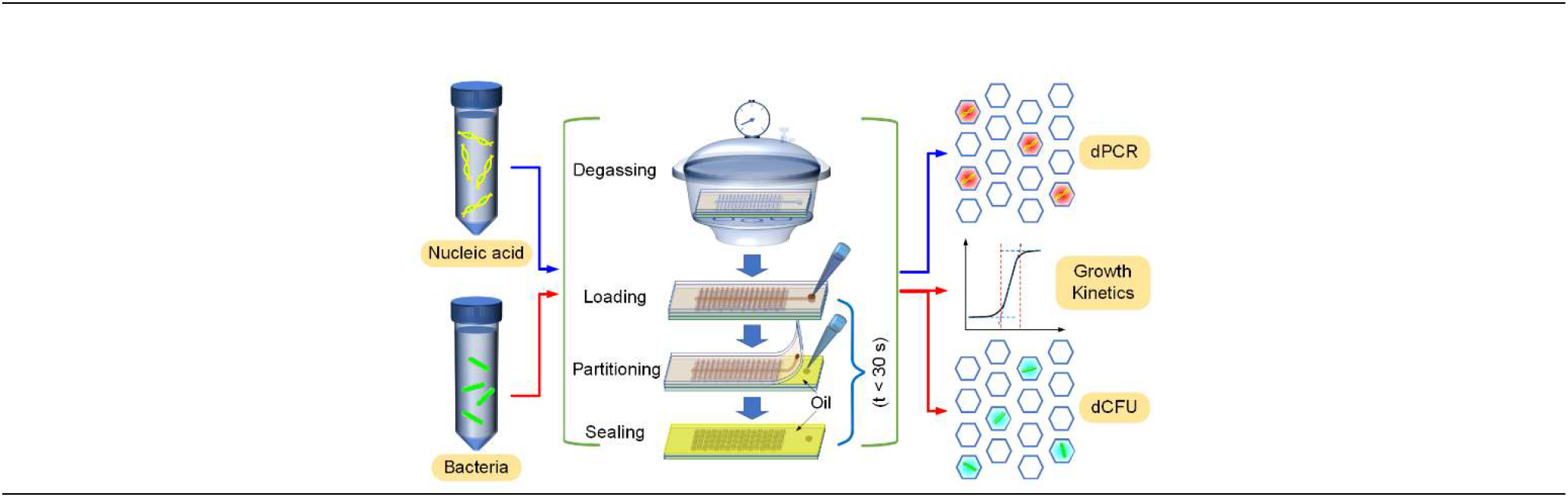

## Supplementary information

### S1. Bacterial Sample Preparation

Revival of a stored *E. coli* BL21 strain was carried out by scraping off splinters of a frozen glycerol stock with a sterile loop and then streaking these splinters onto a LB agar plate for single colonies. After inoculation, the plate was incubated at 37°C until colonies become visible. Next, an isolated colony from the plate was transferred to 5 mL LB broth and incubated for 6 h at 37°C with a shaking rate of 220 rpm.

### S2. Measurement of Bacterial Stock Concentrations via Plating

The *E. coli* BL21 stock sample was diluted with LB broth down to 3.125×10 CFU/mL, 1.250×10^2^ CFU/mL, 5.000×10^2^ CFU/mL. One-hundred μL from each of the 3 dilutions was plated in a LB agar plate, resulting in three plates with expected CFU counts of 3, 12, and 50, respectively. All LB agar plates were incubated at 37 °C for 12 h. After incubation, the number of colonies on each plate was manually counted and recorded directly on the plate. Each measurement was repeated for 3 separate samples, resulting in a total of 9 data points (*i.e.*, 3 titrations in triplicate). The counted number of colonies was plotted against the expected number of colonies, and a linear fit was performed. The linearity was examined and the slope was calculated. Finally, if necessary, the slope of the linear fit line was used to recalculate the bacterial sample concentration. According to the fit line, the concentration of this E. coli stock is recalculated as 6.0 × 10^9^ CFU/mL.

**Figure S1.**
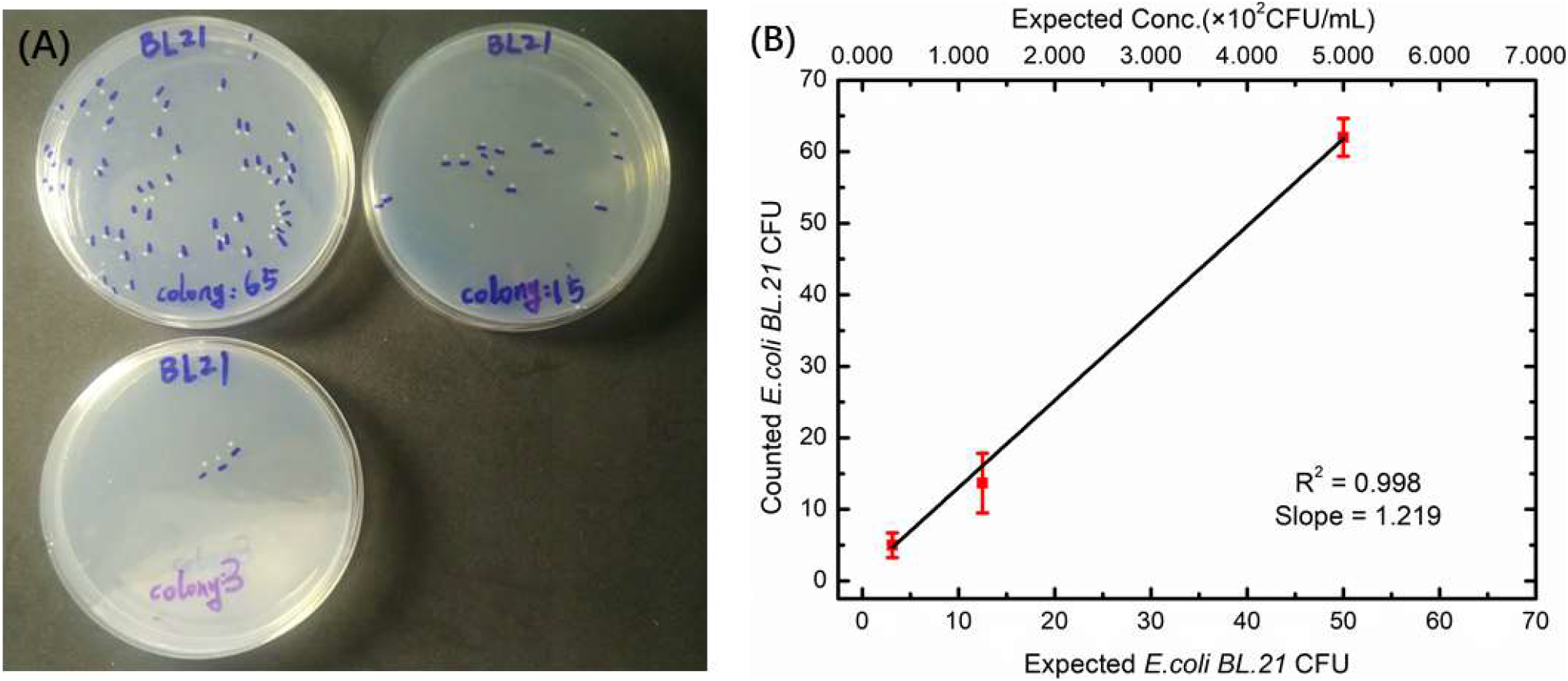
Measurement of *E. coli BL21* stock concentration *via* plating. **(A)** An aliquot of a ~6.0 × 10^9^ CFU/mL *E. coli* stock (originally estimated *via* a single plate) is divided into 3 titrations (3.125 × 10, 1.250 × 10^2^, and 5.000 × 10^2^ CFU/mL), and 100 μL of each titration is plated in LB agar plates, resulting in expected CFU counts of 3, 12, and 50, respectively. After 12-h, 37 °C incubation, the number of colonies from each titration is manually counted and recorded directly on the plate. **(B)** After repeating the same experiment for 3 separate aliquots (*i.e.*, 3 titrations in triplicate), the counted number of colonies is plotted against the expected number of colonies. The slope of the linear fit line is 1.219. Consequently, the concentration of this *E. coli* stock is recalculated as 6.0 × 10^9^ CFU/mL. Error bars represent standard deviations from triplicated experiments.

### S3. Measurement of Fluorescence Intensity of Microcolonies with ImageJ

For determination of the number of viable bacteria in microcolonies, we used ImageJ as tool for measuring the total fluorescence intensity of each microcolony, which is directly proportional to the number of viable bacteria contained in the microcolony. The detailed procedure is shown below:

1. Load an image which is needed to be analyzed. To do this, select: *File > open.*
2. Convert the image to grayscale. To do this, select: *Image > Type > 8-bit Grayscale.*
3. Select a region of interest (ROI) using wand tool. To do this, click on the wand tool in the tool bar and click on a fluorescence region. To achieve a precise selection, double click on the wand tool and adjust the tolerance.
4. Set desired parameters by going to *Analyze > Set Measurements.* Make sure “Area” and “Integrated Density” are checked.
5. Get the value of fluorescence intensity of ROI. To do this, select: *Analyze > Measure.* A window will pop up with the measured values. Copy data into a spreadsheet
6. Repeat for several more microcolonies.

**Figure S2.**
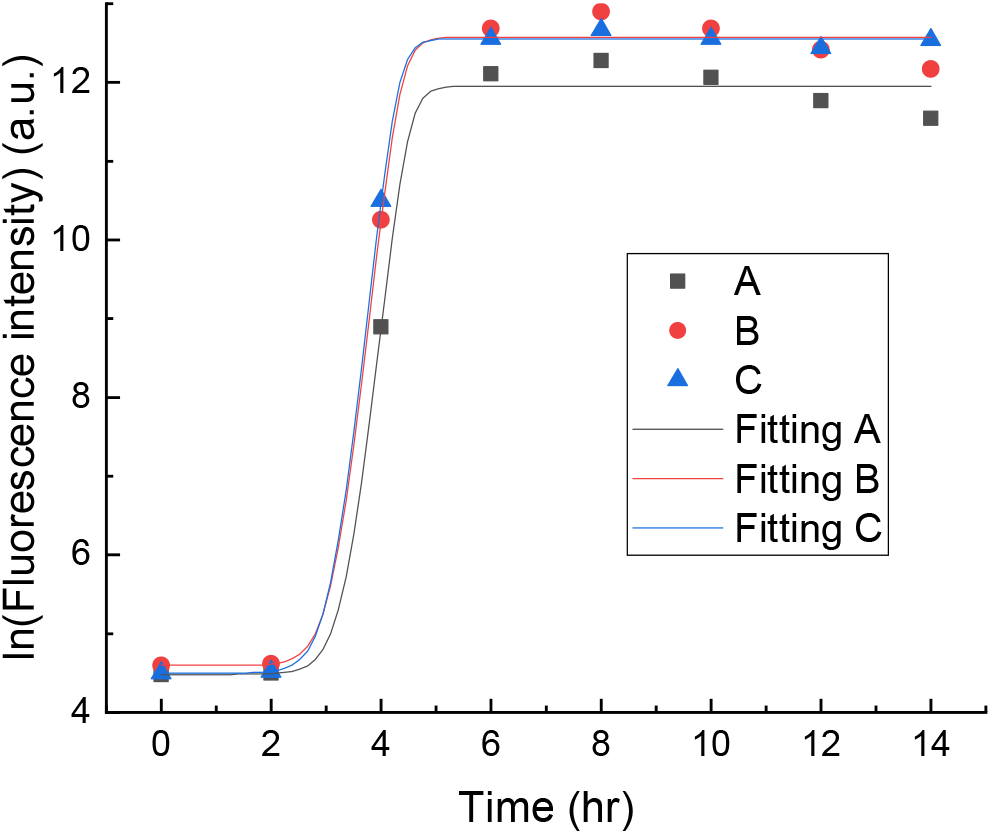
Growth curves produced by fitting the experimental data to Huang primary model. A, B, and C correspond to three different microcolonies labelled in Figure 6a, respectively.

## Notes

### Competing Interest Statement

The authors have declared no competing interest.

